# High-level cognition is supported by information-rich but compressible brain activity patterns

**DOI:** 10.1101/2023.03.17.533152

**Authors:** Lucy L. W. Owen, Jeremy R. Manning

## Abstract

Brain activity patterns are highly flexible and often complex, but also highly structured. Here we examined how fundamental properties of brain activity patterns relate to ongoing cognitive processes. To this end, we applied dimensionality reduction algorithms and pattern classifiers to functional neuroimaging data collected as participants listened to a story, temporally scrambled versions of the story, or underwent a resting state scanning session. These experimental conditions were intended to require di*ff*erent depths of processing and inspire di*ff*erent levels of cognitive engagement. We considered two primary aspects of the data. First, we treated the maximum achievable decoding accuracy across participants as an indicator of the “informativeness” of the recorded patterns. Second, we treated the number of features (components) required to achieve a threshold decoding accuracy as a proxy for the “compressibility” of the neural patterns (where fewer components indicate greater compression). Overall, we found that the peak decoding accuracy (achievable without restricting the numbers of features) was highest in the intact (unscrambled) story listening condition. However, the number of features required to achieve comparable classification accuracy was also lowest in the intact story listening condition. Taken together, our work suggests that our brain networks flexibly reconfigure according to ongoing task demands, and that the activity patterns associated with higher-order cognition and high engagement are both more informative and more compressible than the activity patterns associated with lower-order tasks and lower levels of engagement.

**Significance Statement:** How our brains respond to ongoing experiences depends on what we are doing and thinking about, among other factors. To study how brain activity reflects ongoing cognition, we examined two fundamental aspects of brain activity under di*ff*erent cognitive circumstances: informativeness and compressibility. Informativeness refers to the extent to which brain patterns are both temporally specific and consistent across di*ff*erent people. Compressibility refers to how robust the informativeness of brain patterns is to dimensionality reduction. Brain activity evoked by higher-level cognitive tasks are both more informative *and* more compressible than activity evoked by lower-level tasks. Our findings suggest that our brains flexibly reconfigure themselves to optimize di*ff*erent aspects of how they function according to ongoing cognitive demands.

## Introduction

Large-scale networks, including the human brain, may be conceptualized as occupying one or more positions along on a continuum. At one extreme, every node is fully independent from every other node. At the other extreme, all nodes behave identically. Each extreme optimizes key properties of how the network functions. When every node is independent, the network is maximally *expressive*: if we define the network’s “state” as the activity pattern across its nodes, then every state is equally reachable by a network with fully independent nodes. On the other hand, a network of identically behaved nodes optimizes *robustness*: any subset of nodes may be removed from the network without any loss of function or expressive power, as long as any single node remains. In addition to considering flexibility across space (nodes), these properties may also vary, largely independently, across time. A network is maximally expressive when its nodes’ activity patterns vary in meaningful ways from moment to moment, whereas it is maximally robust to signal corruption when its activity is constant over time. Presumably, most natural systems tend to occupy positions between these temporal and spatial extremes. Under di*ff*erent circumstances, it may even prove benefitial for systems to make di*ff*erent tradeo*ff*s between expressiveness and robustness along the temporal and spatial dimensions. We wondered: might the human brain reconfigure itself to be more flexible or more robust according to ongoing demands? In other words, might the brain reconfigure its connections or behaviors under di*ff*erent circumstances to change its position along these continuums?

Closely related to the above notions of expressiveness versus robustness are measures of how much *information* is contained in a given signal or pattern, and how *redundant* a signal is (Shannon, 1948). Formally, information is defined as the amount of uncertainty about a given variables’ outcomes (i.e., entropy), measured in *bits*, or the optimal number of yes*/*no questions needed to reduce uncertainty about the variable’s outcomes to zero. Highly complex systems with many degrees of freedom (i.e., high flexibility and expressiveness), are more information-rich than simpler or more constrained systems. The redundancy of a signal denotes the di*ff*erence between how expressive the signal *could* be (i.e., proportional to the number of unique states or symbols used to transmit the signal) and the actual information rate (i.e., the entropy of each individual state or symbol). If a brain network’s nodes are fully independent, then the number of bits required to express a single activity pattern is proportional to the number of nodes. The network would also be minimally redundant, since the status of every node would be needed to fully express a single brain activity pattern. If a brain network’s nodes are fully coupled and identical, then the number of bits required to express a single activity pattern is proportional to the number of unique states or values any individual node can take on. Such a network would be highly redundant, since knowing any individual node’s state would be su*ffi*cient to recover the full-brain activity pattern. Highly redundant systems are also robust, since there is little total information loss due to removing any given observation.

We take as a given that brain activity is highly flexible: our brains can exhibit nearly infinite varieties of activity patterns. This flexibility implies that our brains’ activity patterns are highly information rich. However, brain activity patterns are also highly structured. For example, full-brain correlation matrices are stable within (Finn et al., 2017, 2015; Gratton et al., 2018) and across (Cole et al., 2014; Glerean et al., 2012; Gratton et al., 2018; Yeo et al., 2011) individuals. This stability suggests that our brains’ activity patterns are at least partially constrained, for example by anatomical, external, or internal factors. Constraints on brain activity that limit its flexibility decrease expressiveness (i.e., its information rate). However, constraints on brain activity also increase its robustness to noise (e.g., “missing” or corrupted signals may be partially recovered). For example, recent work has shown that full-brain activity patterns may be reliably recovered from only a relatively small number of implanted electrodes (Owen et al., 2020; Scangos et al., 2021). This robustness property suggests that the relevant signal (e.g., underlying factors that have some influence over brain activity patterns) are compressible.

To the extent that brain activity patterns contain rich task-relevant information, we should be able to use the activity patterns to accurately di*ff*erentiate between di*ff*erent aspects of a task (e.g., using pattern classifiers; Norman et al., 2006). For example, prior work has shown a direct correspondence between classification accuracy and the information content of a signal (Alvarez, 2002). To the extent that brain activity patterns are compressible, we should be able to generate simplified (e.g., lower dimensional) representations of the data while still preserving the relevant or important aspects of the original signal. In general, information content and compressibility are often related but are also dissociable (Fig. 1). If a given signal (e.g., a representation of brain activity patterns) contains more information about ongoing cognitive processes, then the peak decoding accuracy should be high. In the simulations shown in Figure 1C we construct synthetic datasets that have high or low levels of informativeness by varying temporal autocorrelations in the data (see *Synthetic data*). If a signal is compressible, then a low-dimensional embedding of the signal will be similarly informative as the original signal. In the simulations shown in Figure 1C we construct synthetic datasets that have high or low levels of compressibility by varying the covariance structure across features (see *Synthetic data*). As shown in Figure 1D, highly informative datasets yield higher decoding accuracies than less informative datasets (i.e., the peaks of the curves in the top panels of Fig. 1D are higher than the peaks of the curves in the bottom panels). Highly compressible datasets show steeper slopes when we plot decoding accuracy as a function of the number of components used to represent the data (i.e., the slopes of the curves in the left panels of Fig. 1D are steeper than the slopes of the curves in the right panels). Whereas characterizing the informativeness and compressibility of synthetic data can be instructive, we are ultimately interested in understanding how these properties relate to brain activity patterns recorded under di*ff*erent cognitive circumstances.

**Figure 1:**
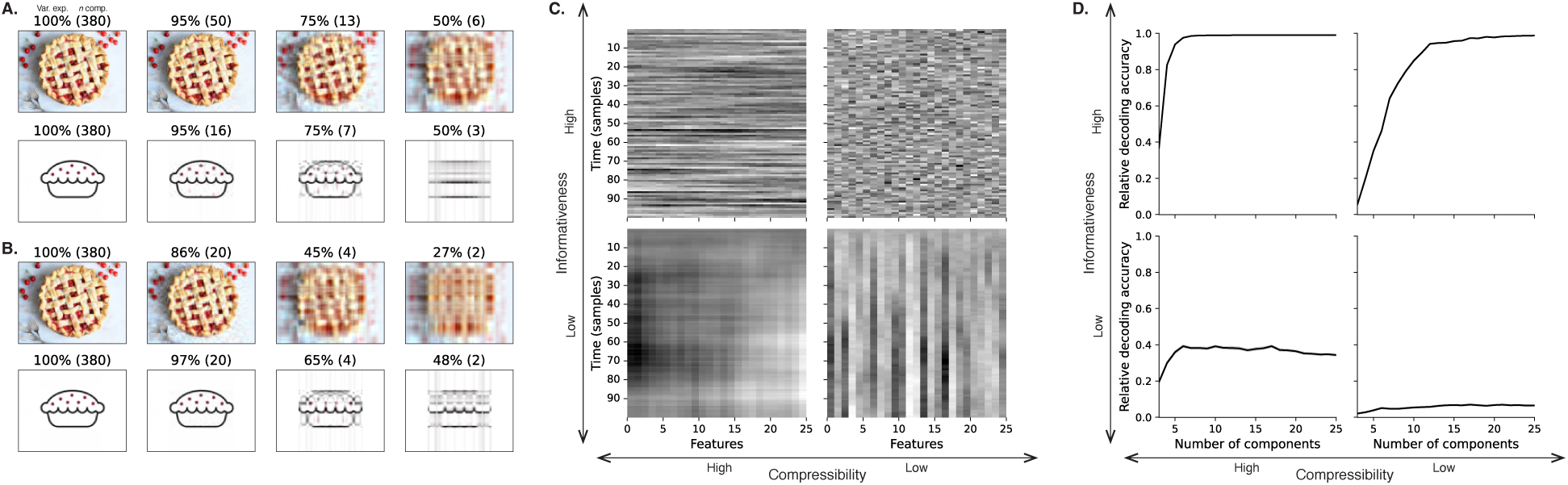
Information content and compressibility. **A. Variance explained for two images.** Consider two example images: a photograph and simple drawing. The images have the same resolutions, but their content is very di*ff*erent. The photograph is more information rich: when we compare the original photograph and drawing (leftmost column), we can see that the photograph captures much more detail. We can also apply principal components analysis to the images, treating the rows of the images as “observations.” Across columns, we identified the numbers of components required to explain 100%, 95%, 75%, or 50% of the cumulative variance in each image (the 100% columns denote the original images). The numbers of components are indicated in parentheses, and the resulting “compressed” images are displayed. This analysis reveals that the drawing is more compressible: just 16 components can explain 95% of the variance in the drawing, whereas 50 components are required to explain 95% of the variance in the photograph. **B. Representing two images with di**ff**erent numbers of components.** Using the same principal component decompositions as in Panel A, we computed the cumulative proportion of variance explained with 380 (original images), 20, 4, or 2 components. This analysis provides another way of characterizing the compressibility of each image by showing that the same number of components can explain more variance in the drawing than in the photograph. **C. Template data from four synthetic datasets.** We constructed four synthetic datasets, each comprising 25 features (columns) observed across 100 samples (rows) from each of 10 simulated “participants.” The datasets were constructed to contain di*ff*erent levels of informativeness and compressibility (see *Synthetic data*). **D. Decoding accuracy by number of components.** For each synthetic dataset, we trained across-participant classifiers to decode timepoint labels. Each panel displays the decoding accuracy as a function of the number of components used to represent the data. Error ribbons denote bootstrap-estimated 95% confidence intervals.

Several recent studies suggest that the complexity of brain activity is task-dependent, whereby simpler tasks with lower cognitive demands are reflected by simpler and more compressible brain activity patterns, and more complex tasks with higher cognitive demands are reflected by more complex and less compressible brain activity patterns (Mack et al., 2020; Owen et al., 2021). These patterns hold even when the stimulus itself is held constant (Mack et al., 2020). These findings complement other work suggesting that functional connectivity (correlation) patterns are task-dependent (Cole et al., 2014; Finn et al., 2017; Owen et al., 2020), although see Gratton et al. (2018). Higher-order cognitive processing of a common stimulus also appears to drive more stereotyped task-related activity and functional connectivity across individuals (Hasson et al., 2008; Lerner et al., 2011; Simony & Chang, 2020; Simony et al., 2016).

The above studies are consistent with two potential descriptions of how cognitive processes are reflected in brain activity patterns. One possibility is that the information rate of brain activity increases during more complex or higher-level cognitive processing. If so, then the ability to reliably decode cognitive states from brain activity patterns should improve with task complexity or with the level (or “depth”) of cognitive processing. A second possibility is that the compressibility of brain activity patterns decreases during more complex or higher-level cognitive processing. If so, then individual features of brain recordings should carry more information (over and above the information carried by other features) during complex or high-level (versus simple or low-level) cognitive tasks. The tradeo*ff*s between these two aspects of brain activity may also vary across brain regions or networks, for example according to each region’s functional role.

We used a previously collected neuroimaging dataset to estimate the extent to which each of these two possibilities might hold. The dataset we examined comprised functional magnetic resonance imaging (fMRI) data collected as participants listened to an audio recording of a 7-minute story, temporally scrambled recordings of the story, or underwent a resting state scan (Simony et al., 2016). Each of these experimental conditions evokes di*ff*erent depths of cognitive processing (Hasson et al., 2008; Lerner et al., 2011; Owen et al., 2021; Simony et al., 2016). We used across-participant classifiers to decode listening times in each condition, as a proxy for how “informative” the task-specific activity patterns were (Simony & Chang, 2020). We also used principle components analysis to generate lower-dimensional representations of the activity patterns. We then repeated the classification analyses after preserving di*ff*erent numbers of components and examined how classification accuracy changed across the di*ff*erent experimental conditions.

## Results

We sought to understand whether higher-level cognition is reflected by more reliable and informative brain activity patterns, and how compressibility of brain activity patterns relates to cognitive complexity. We developed a computational framework for systematically assessing the informativeness and compressibility of brain activity patterns recorded under di*ff*erent cognitive circumstances. We used across-participant decoding accuracy (see *Forward inference and decoding accuracy*) as a proxy for informativeness. To estimate the compressibility of the brain patterns, we used group principal components analysis (PCA) to project the brain patterns into *k*-dimensional spaces, for di*ff*erent values of *k* (see *Hierarchical Topographic Factor Analysis (HTFA)* and *Principal components analysis (PCA)*). For more compressible brain patterns, decoding accuracy should be more robust to small values of *k*.

We analyzed a dataset collected by Simony et al. (2016) that comprised four experimental conditions. These conditions exposed participants to stimuli that systematically varied in cognitive engagement. In the *intact* experimental condition, participants listened to an audio recording of a 7-minute Moth Radio Hour story, *Pie Man*, by Jim O’Grady. In the *paragraph*-scrambled experimental condition, participants listened to a temporally scrambled version of the story, where the paragraphs occurred out of order, but where the same set of paragraphs was presented over the entire listening interval. All participants in this condition experienced the scrambled paragraphs in the same order. In the *word*-scrambled experimental condition, participants listened to a temporally scrambled version of the story, where the words occurred in a random order. Again, all participants in this condition experienced the scrambled words in the same order. Finally, in the *rest* experimental condition, participants lay in the scanner with no overt stimulus, while keeping their eyes open and blinking as needed. This public dataset provided a convenient means for testing our hypothesis that di*ff*erent levels of cognitive processing and engagement a*ff*ect how informative and compressible the associated brain patterns are.

To evaluate the relation between informativeness and compressibility for brain activity from each experimental condition, we trained a series of across-participant temporal decoders on compressed representations of the data. Figure 2A displays the decoding accuracy as a function of the number of principal components used to represent the data (also see Fig. S1). Several patterns were revealed by the analysis. First, in general (i.e., across experimental conditions), decoding accuracy tends to improve as the number of components are increased. However, decoding accuracy peaked at higher levels for experimental conditions that exposed participants to cognitively richer stimuli (Fig. 2D). The peak decoding accuracy was highest for the “intact” condition (versus paragraph: *t*(99) *=* 35.205, *p* < 0.001; versus word: *t*(99) *=* 43.172, *p* < 0.001; versus rest: *t*(99) *=* 81.361, *p* < 0.001), next highest for the “paragraph” condition (versus word: *t*(99) *=* 6.243, *p* < 0.001; versus rest: *t*(99) *=* 50.748, *p* < 0.001), and next highest for the “word” condition (versus rest: *t*(99) *=* 48.791, *p* < 0.001). This ordering implies that cognitively richer conditions evoke more stable brain activity patterns across people.

**Figure 2:**
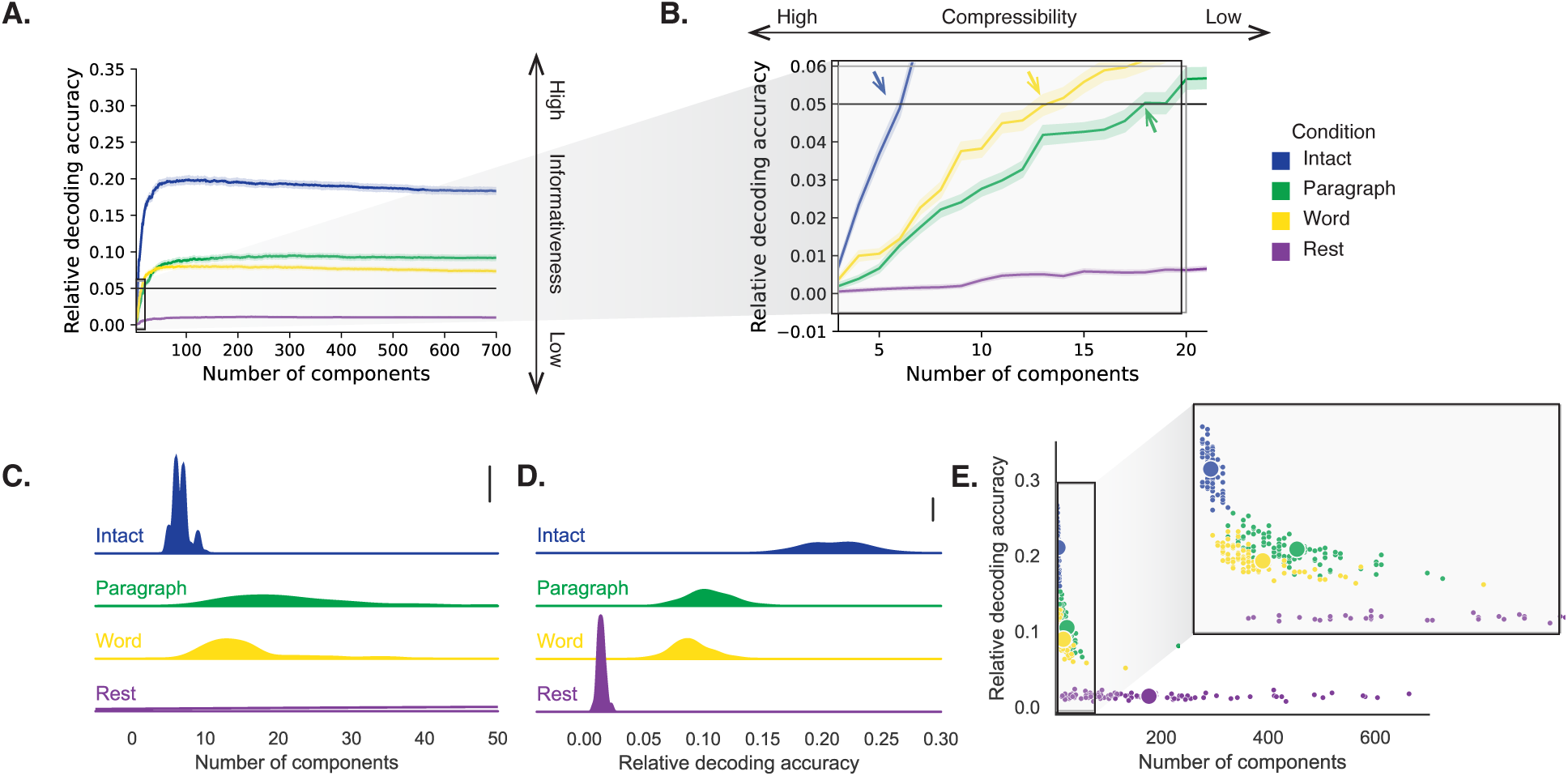
Decoding accuracy and compression. **A. Decoding accuracy by number of components.** Ribbons of each color display cross-validated decoding performance for each condition (intact, paragraph, word, and rest), as a function of the number of components (features) used to represent the data. For each condition, decoding accuracy has been normalized by subtracting expected “chance” decoding performance (estimated as <u>^1^</u>, where *T* is the number of timepoints in the given condition). This normalization was intended to enable fairer comparisons across conditions; a relative decoding accuracy of 0.0 denotes “chance” performance. The horizontal black line denotes 5% decoding accuracy (used as a reference in Panel B). **B. Numbers of components required to reach a fixed decoding accuracy threshold, by condition.** The panel displays a zoomed-in view of the inset in Panel A. Intersections between each condition’s decoding accuracy curve and the 5% decoding accuracy reference line are marked by arrows. All error ribbons in Panels A and B denote bootstrap-estimated 95% confidence intervals. **C. Estimating “compressibility” for each condition.** The probability density plots display the numbers of components required to reach at least 5% (corrected) decoding accuracy, across all cross validation runs. For the “rest” condition, where decoding accuracy never reached 5% accuracy, we display the distribution of the numbers of components required to achieve the peak decoding accuracy in each fold. (This distribution is nearly flat, as also illustrated in Panel E.) The scale bar denotes a height of 0.01. **D. Estimating “informativeness” for each condition.** The probability density plots display the peak (corrected) decoding accuracies achieved in each condition, across all cross validation runs. The scale bar denotes a height of 0.01. **E. Informativeness versus compressibility.** Each dot displays the minimum number of components required to achieve at least 5% corrected decoding accuracy (*x*-coordinate; for the rest condition the numbers of components are estimated using the peak decoding accuracies as in Panel C) and the peak (corrected) decoding accuracy (*y*-coordinate). The smaller dots correspond to individual cross validation runs and the larger dots display the averages across runs. The inset displays a zoomed-in view of the indicated region in the main panel.

The cognitively richer conditions also displayed steeper initial slopes. For example, the intact condition decoders reached an arbitrarily chosen threshold of 5% accuracy using fewer components than the paragraph condition decoders (*t*(99) *=* −7.429, *p* < 0.001) or word condition decoders (*t*(99) *=* −7.300, *p* < 0.001), and decoding accuracy never exceeded 5% for the rest condition. This suggests that brain activity patterns evoked by cognitively richer conditions are more compressible, such that representing the data using the same number of principal components provides more information to the temporal decoders (Figs. 2B, C). Taken together, as shown in Figure 2E, we found that brain activity patterns evoked by cognitively richer conditions tended to be both more informative (i.e., associated with higher peak decoding accuracies) *and* more compressible (i.e., requiring fewer components to achieve the 5% accuracy threshold).

If informativeness (to the temporal decoders) and compressibility vary with the cognitive richness of the stimulus, might these measures also vary over time *within* a given condition? For example, participants in the intact condition might process the ongoing story more deeply later on in the story (compared with earlier in the story) given the additional narrative background and context they had been exposed to by that point. To examine this possibility, we divided each condition into four successive time segments. We computed decoding curves (Fig. 3A) and the numbers of components required to achieve 5% decoding accuracy (Fig. 3B) for each segment and condition. We found that, in the two most cognitively rich conditions (intact and paragraph), both decoding accuracy and compressibility, as reflected by the change in decoding curves, increased with listening time (e.g., at the annotated reference point of *k =* 20 components in Fig. 3C: intact: *t*(99) *=* 7.915, *p* < 0.001; paragraph: *t*(99) *=* 2.354, *p =* 0.021). These changes may reflect an increase in comprehension or depth of processing with listening time. In contrast, the decoding accuracy and compressibility *decreased* with listening time in the word condition (*t*(99) *=* −10.747, *p* < 0.001) and rest condition (*t*(99) *=* −22.081, *p* < 0.001). This might reflect the depletion of attentional resources in the less-engaging word and rest conditions.

**Figure 3:**
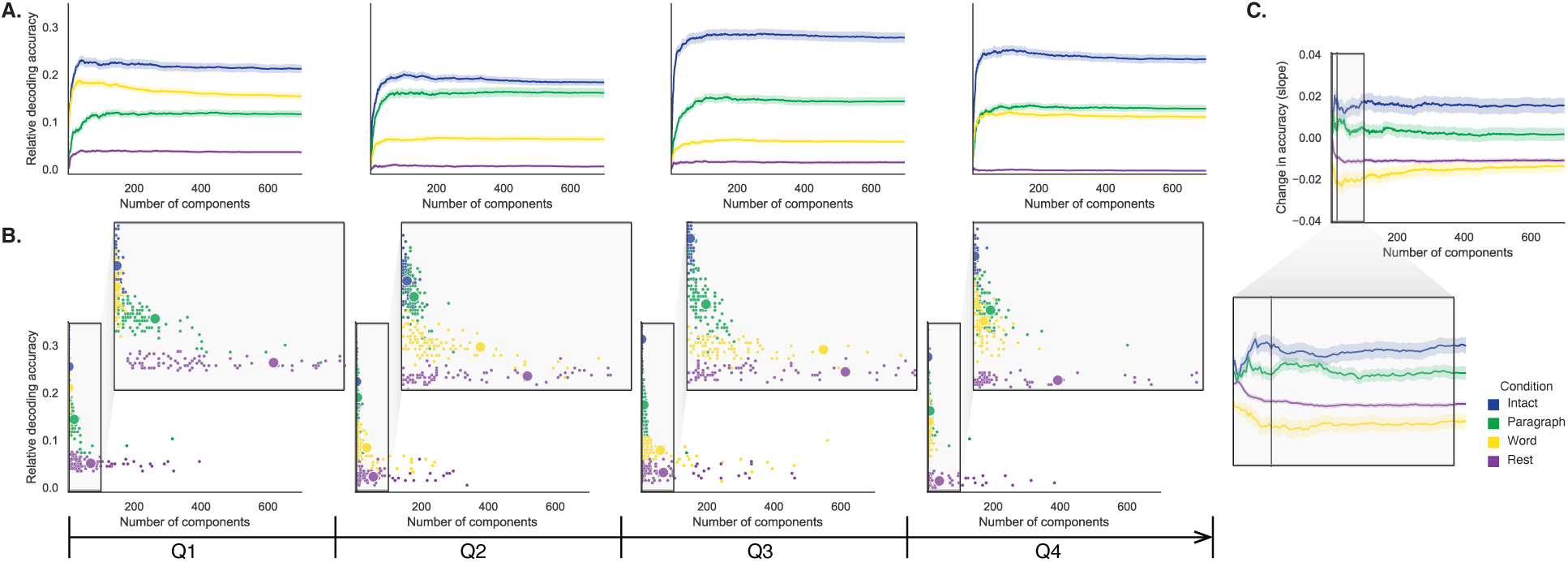
Changes in decoding accuracy and compression over time. **A. Decoding accuracy by number of components, by story segment.** Each family of curves is plotted in the same format as Figure 2A but reflects data only from one quarter of the dataset. **B. Informativeness versus compressibility by condition and segment.** Each scatter plot is in the same format as Figure 2E, but reflects data only from one quarter of the dataset. **C. Change in decoding accuracy over time, by number of components.** For each number of components (*x*-axis) and condition (color), we fit a regression line to the decoding accuracies obtained for the first, second, third, and fourth quarters of the dataset (corresponding to the columns of Panels A and B). The *y*-axis denotes the slopes of the regression lines. The black vertical line marks *k =* 20 components, as referenced in the main text. All error ribbons denote bootstrap-estimated 95% confidence intervals.

These results make some intuitive sense. As the contextual information available to participants increases (i.e., over time in the cognitively rich intact and paragraph conditions), it makes sense that this might constrain neural responses to a greater extent. While this pattern may not necessarily hold for *every* possible story or stimulus, we suspect that it is generally the case that our knowledge about what is happening in a story tends to increase as we experience more of it. In turn, this could lead to greater consistency in di*ff*erent people’s interpretations of and neural responses to the stimulus. Similarly, as participants are left to “mind wander,” or as they experience mental fatigue (i.e., over time in the less cognitively rich word and rest conditions), we suggest that this might lead to greater variability in neural responses across people, resulting in lower decoding accuracy. Again, it is not necessarily the case that every possible “unengaging” stimulus will lead to greater neural variability as time progresses, but we suspect this phenomenon is likely to hold for a variety of such stimuli. These findings replicate at least to a limited extent (e.g., across both cognitively rich conditions and across both cognitively impoverished conditions, and for the di*ff*erent groups of participants in each of those conditions). However, determining whether these patterns generalize to other stimuli would require additional study (with new stimuli).

If the informativeness and compressibility of brain activity patterns vary over time, might these properties also vary across brain networks? We used a network parcellation identified by Yeo et al. (2011) to segment the brain into seven distinct networks. The networks can be sorted (roughly) in order from lower-level to higher-level cortex as follows (Figs. 4A–C): visual, somatomotor, dorsal attention, ventral attention, limbic, frontoparietal, and default mode. Next, we computed decoding curves separately for the activity patterns within each network and identified each network’s inflection point, for each experimental condition. Moving from low-order networks to higher-order networks, we found that decoding accuracy tended to increase in the higher-level experimental conditions and decrease (slightly) in the lower-level experimental conditions (Fig. 4D, E; Spearman’s rank correlation between decoding accuracy and network order: intact: ρ *=* 0.362, *p* < 0.001; paragraph: ρ *=* 0.441, *p* < 0.001; word: ρ *=* −0.102, *p =* 0.007; rest: ρ *=* −0.354, *p* < 0.001). This suggests that higher-order networks may carry more content-relevant or stimulus-driven “information.” We found no clear trends in the proportions of components required to achieve 5% decoding accuracy across networks or conditions (Fig. 4F). We note that the limbic network we considered here often overlaps with low (imaging) signal regions, and therefore it may be di*ffi*cult to draw strong conclusions about this network’s informativeness or compressibility. We also considered the possibility that the correlations with network order might be influenced by the numbers of nodes in each network. We designed a permutation-based procedure to address this possibility, whereby we repeated the above analyses using shu*ffl*ed network labels (see *Network permutation tests*). The correlations between decoding accuracy and network order were reliably more positive than the shu*ffl*ed correlations for the intact (*t*(1998) *=* 276.431, *p* < 0.001) and paragraph (*t*(1998) *=* 330.334, *p* < 0.001) conditions, and reliably more negative for the word (*t*(1998) *=* −16.386, *p* < 0.001) and rest (*t*(1998) *=* −318.631, *p* < 0.001) conditions. These results suggest that the correlations between decoding accuracy and network order were not driven solely by the numbers of nodes in each network.

**Figure 4:**
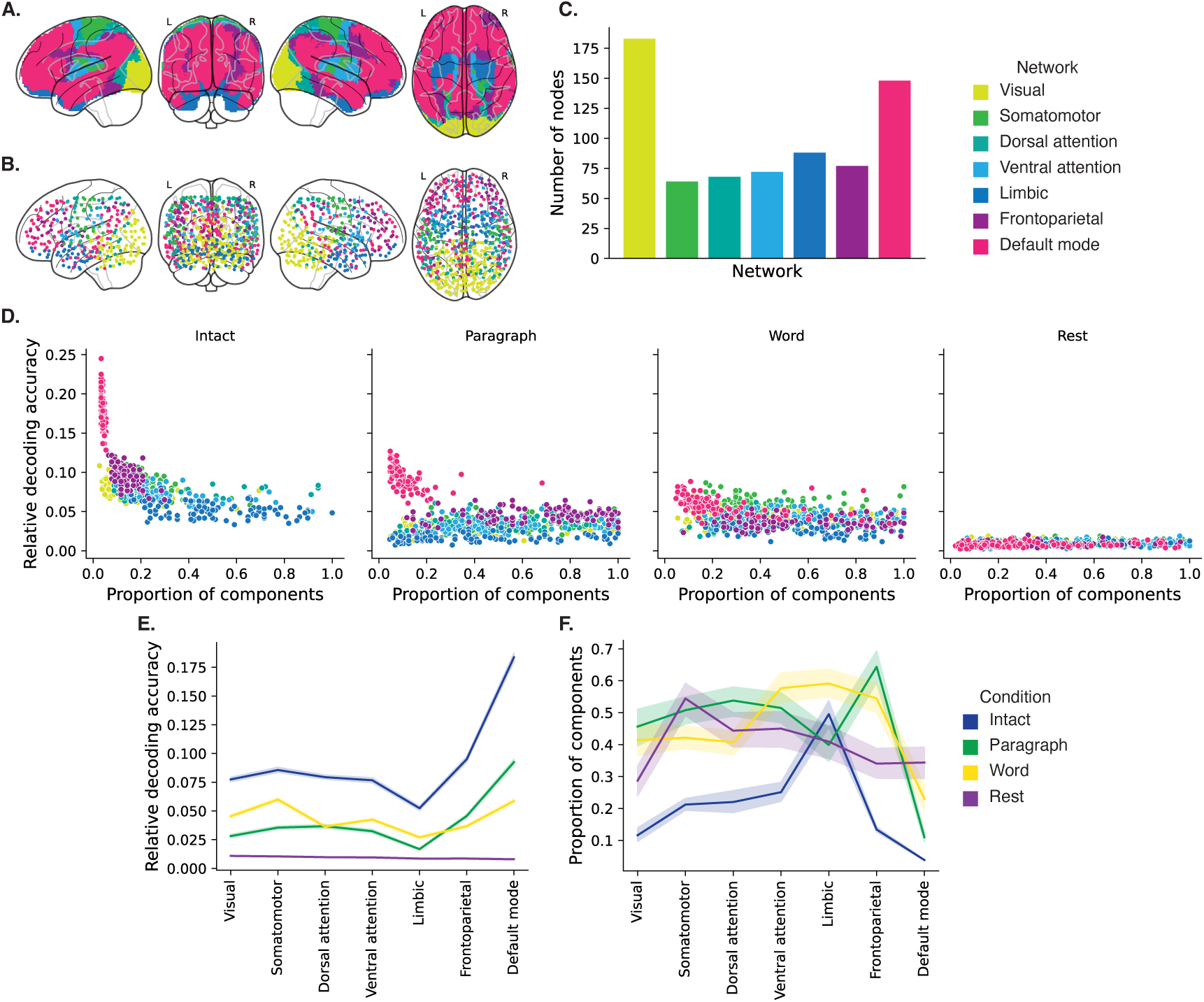
Network-specific decoding accuracy and compression. **A. Network parcellation map.** The glass brain displays the location of each of seven networks identified by Yeo et al. (2011). **B. Node labels.** We assigned network labels to each of 700 Hierarchical Topographic Factor Analysis-derived nodes (see *Hierarchical topographic factor analysis (HTFA)*). For each node, we evaluated its radial basis function (RBF) at the locations of every voxel in the parcellation map displayed in Panel A. We summed up these RBF weights separately for the voxels in each network. We defined each node’s network label as its highest-weighted network across all voxels. **C. Per-network node counts.** The bar plot displays the total number of HTFA-derived nodes assigned to each network. **D. Informativeness versus compressibility by network and condition.** Each dot corresponds to a single cross validation run. The dots’ coordinates denote the minimum proportion of components (relative to the number of nodes in the given network) required to achieve at least 5% corrected decoding accuracy (*x*-coordinate; for the rest condition these proportions are estimated using the peak decoding accuracies) and peak (corrected) decoding accuracy (*y*-coordinate). We repeated the analysis separately for each network (color) and experimental condition (column). **E. Informativeness by network.** For each network, sorted (roughly) from lower-order networks on the left to higher-order networks on the right, the curves display the peak (corrected) decoding accuracy for each experimental condition (color). **F. Compressibility by network.** For each network, the curves display the proportion of components (relative to the total possible number of components displayed in Panel C) required to achieve at least 5% decoding accuracy (or, for the rest condition, peak decoding accuracy, as described above). Error ribbons in Panels E and F denote bootstrap-estimated 95% confidence intervals.

Whereas the above analyses examined di*ff*erent networks in isolation, how does full-brain (i.e., potentially multi-network) activity patterns reflected by di*ff*erent principal components vary across di*ff*erent experimental conditions? As shown in Figure 5, we used Neurosynth (Rubin et al., 2017) to identify, for each component, the associations with each of 80 themes (see *Reverse inference*). In general, the first principal components across all of the experimental conditions tended to weigh most heavily on themes related to cognitive control, memory, language processing, attention, and perception. Other components appeared to vary more across conditions.

**Figure 5:**
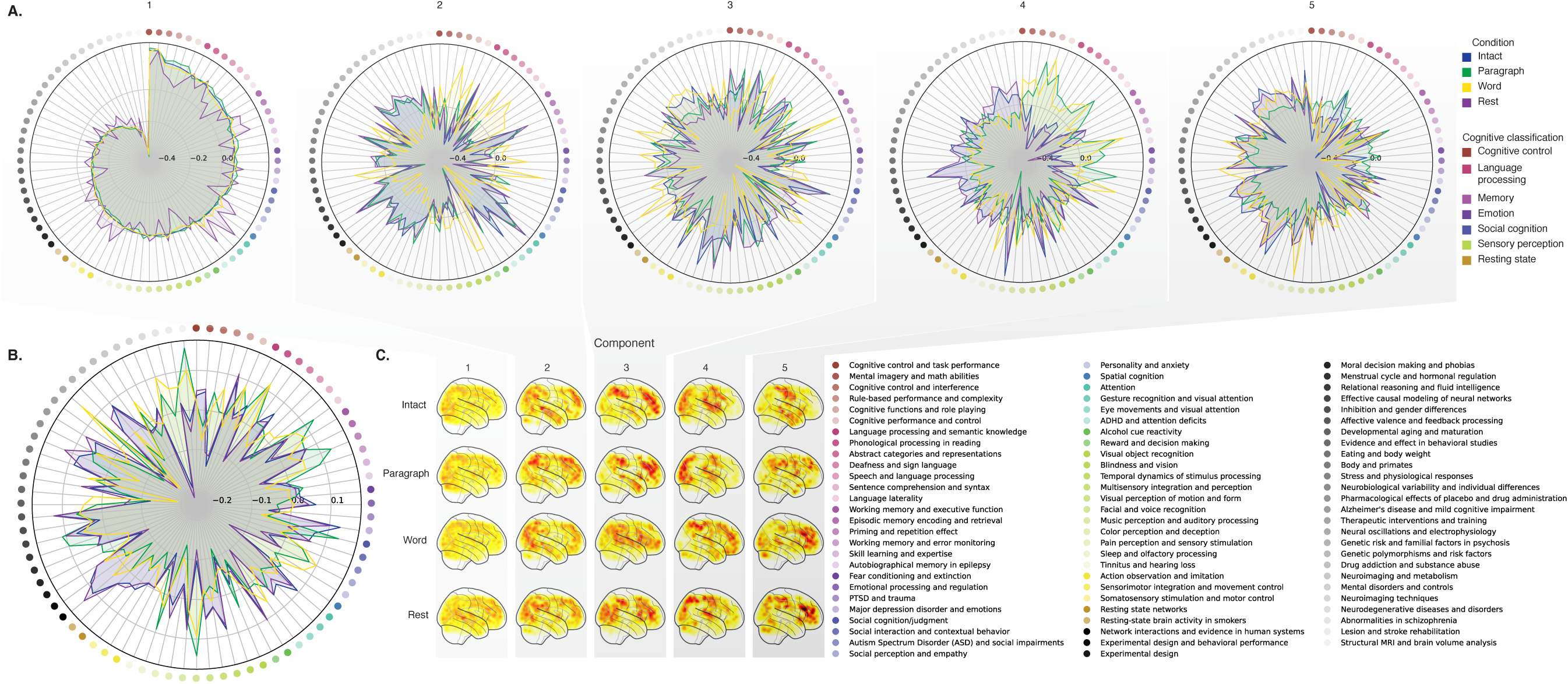
Neurosynth topic weightings by component. We used a reverse inference procedure (see *Reverse inference*) to estimate the correspondences between brain images and a set of 80 topics derived from the Neurosynth database of neuroimaging articles (Rubin et al., 2017). **A. Topic correlations by component.** For each of the top five highest-weighted principal components (columns) derived from each experimental condition (colors), the radar plots display the correlations between the component images and the per-topic images derived for each topic (topic identities are denoted by the blue dots; a legend for the topic labels is in the lower right of the figure). An annotated list of the top-weighted topics for each component and condition may be found in Figure S2 and the top-weighted terms for each topic may be found in Table S1. **B. Topic correlations averaged across components.** The radar plot is in the same format as the plots in Panel A, but here we display the per-condition average correlations across the top five components (for each condition) reflected in Panel A. **C. Component images.** Each plot displays a right sagittal view of a glass brain projection of the top five principal components (columns) for each experimental condition (rows). Additional projections for each component may be found in Figure S3.

To gain further insights into which brain functions might be most closely associated with the top-weighted components from each experimental condition, we manually grouped each Neurosynth-derived topic into a smaller set of core cognitive functions. Separately for each component, we computed the average weightings across all topics that were tagged as being associated with each of these cognitive functions (Figs. 6A, S5A). To help visualize these associations, we used the patterns of associations for each component to construct graphs whose nodes were experimental conditions and cognitive functions (Figs. 6B, S5B). We also computed correlations between the sets of per-topic weightings from each of the top-weighted components from each experimental condition (Fig. 6C, D) and between the brain maps for each condition’s components (Fig. S5C, D). Taken together, we found that each component appeared to weigh on a fundamental set of cognitive functions that varied by experimental condition. For example, the top principal components from every condition weighed similarly (across conditions) on the full set of Neurosynth topics (Fig. 5A) and cognitive functions (Figs. 6A, B and S5 A, B), suggesting that these components might reflect a set of functions or activity patterns that are common across all conditions. The second components’ weightings were similar across the intact, paragraph, and rest conditions (highest-weighted functions: cognitive control, memory, social cognition, and resting state), but di*ff*erent for the word condition (highest-weighted functions: sensory perception and cognitive control). The fourth components’ weighting grouped the paragraph and word conditions (highest-weighted functions: memory, language processing, and cognitive control) and the intact and rest conditions (highest-weighted functions: emotion, social cognition). We also used ChatGPT (OpenAI, 2023) to sort the list of manually tagged cognitive functions from lowest-level to highest-level (Tab. S2, Fig. 6E; also see *Ranking cognitive processes*). We found that higher-level functions tended to be weighted more heavily by top components from the intact and paragraph conditions than lower-level functions (intact vs. word: *t*(198) *=* 11.059, *p* < 0.001; intact vs. rest: *t*(198) *=* 3.699, *p* < 0.001; paragraph vs. word: *t*(198) *=* 13.504, *p* < 0.001; paragraph vs. rest: *t*(198) *=* 4.812, *p* < 0.001; also see *Ranking cognitive processes*). The top components from the word condition showed the opposite tendency, whereby *lower*-level functions tended to be weighted more heavily than higher-level functions (word vs. rest: *t*(198) *=* −7.315, *p* < 0.001). The weighting trends for the intact and paragraph conditions were not reliably di*ff*erent (*t*(198) *=* −0.479, *p =* 0.633). The components from the rest condition showed only a small trending di*ff*erence in the weights associated with high-level versus low-level functions (rest vs. 0: *t*(99) *=* 1.836, *p =* 0.081). These findings suggest that when participants were engaged more strongly (in the more engaging intact and paragraph conditions), their dominant neural patterns reflected higher-level cognitive functions. In contrast, when participants were engaged less strongly (in the less engaging word and rest conditions), their dominant neural patterns reflected lower-level cognitive functions. Although they were highly statistically reliable, it is also important to note that these latter e*ff*ects are also relatively small (e.g., the slopes for *all* of the experimental conditions are numerically close to zero; Fig. 6E). We suggest that this phenomenon may merit further investigation in future work.

**Figure 6:**
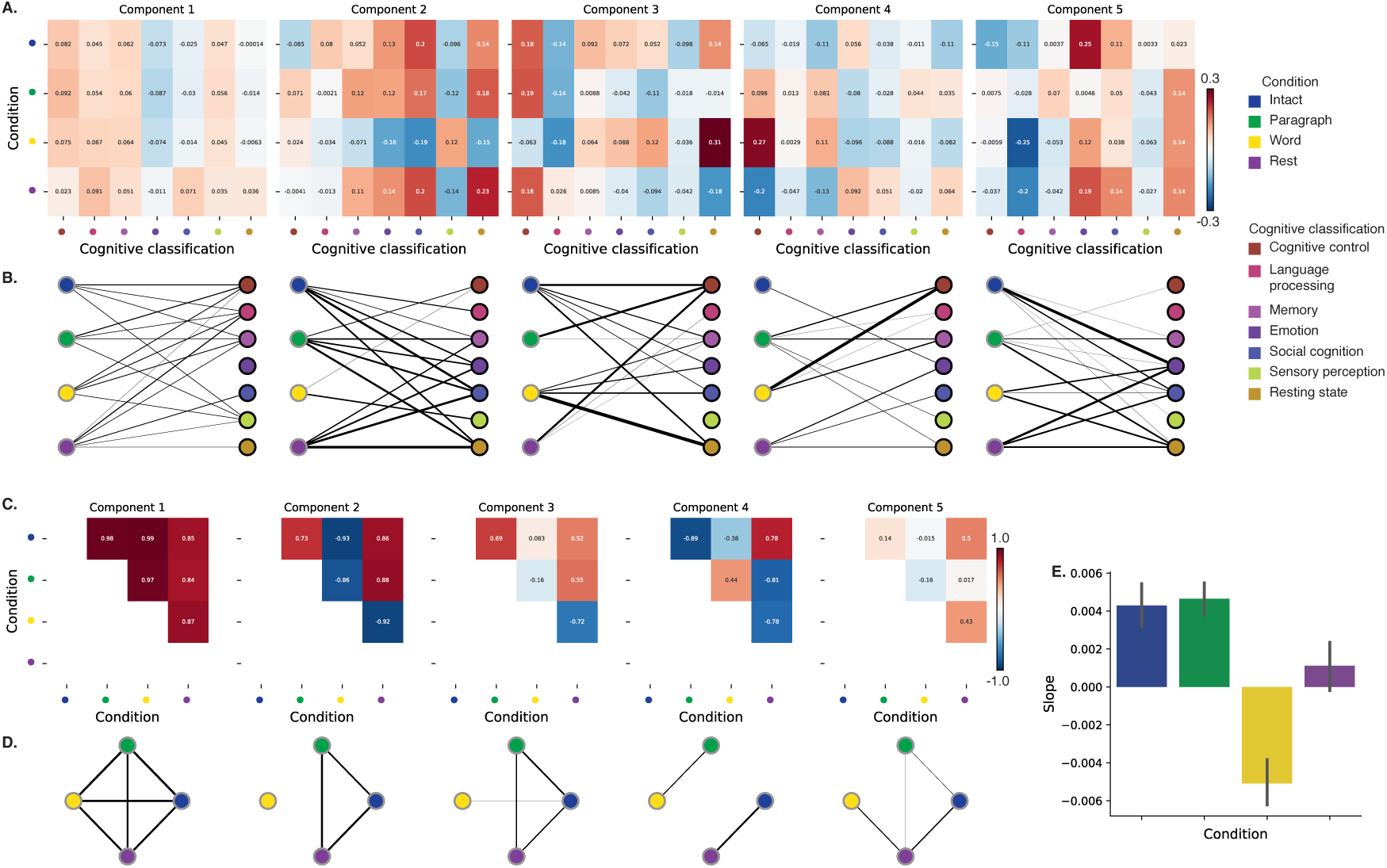
Summary of functions associated with top-weighted components by condition. **A. Top-weighted topics by condition.** Here we display per-condition (rows, indicated by colored dots) topic correlations, averaged across topics that pertain to each of several broad cognitive functions (columns within each sub-panel, indicated by colored dots). Each sub-panel reflects correlations for the components indicated in the panel titles. Table S1 provides a list of each topic’s top-weighted terms, along with each topic’s manually labeled cognitive classification. A full list of the topics most highly associated with each component may be found in Figure S2. **B. Associations between per-condition components and cognitive functions.** The network plots denote positive average correlations between the component images for each condition (gray-outlined dots on the left sides of each network; colors denote conditions) and topic-specific brain maps associated with each indicated cognitive function (black-outlined dots on the right sides of each network; colors denote cognitive functions). The line thicknesses are proportional to the correlation values (correlation coe*ffi*cients are noted in the heat maps in Panel A). **C. Correlations between each principal component, by condition.** The heat maps display the correlations between the brain maps (Fig. S3) for each principal component (sub-panel), across each pair of conditions (rows and columns of each sub-panel’s matrix, indicated by colored dots). **D. Associations between per- condition components, by component.** Each sub-panel’s network plot summarizes the pattern of correlations between the *n*^th^ top-weighted principal components (sub-panel) for each experimental condition (gray-outlined dots). The line thicknesses are proportional to the correlation values (correlation coe*ffi*cients are noted in the heat maps in Panel C). **E. Change in weights by condition.** The bar heights reflect the slopes of regression lines, fit separately to the top five components from each condition, between the ChatGPT-derived “rank” of each cognitive classification and the correlations between the component and topic maps associated with cognitive processes at the given rank (see *Ranking cognitive processes*) Error bars denote bootstrap-estimated 95% confidence intervals. Also see Fig. S5 for additional information.

## Discussion

We examined fMRI data collected as participants listened to an auditory recording of a story, scrambled recordings of the story, or underwent a resting state scan. We found that cognitively richer stimuli evoked more reliable (i.e., consistent across people) and information rich brain activity patterns. The brain patterns evoked by cognitively richer stimuli were also more compressible, in that each individual component provided more “signal” to temporal decoders relative to components of data from less cognitively rich conditions (Fig. 2). Over time (e.g., as the experiment progressed), these phenomena were strengthened. Specifically, across story segments, data from more cognitively rich conditions became more informative and compressible, and data from less cognitively rich conditions became *less* informative and compressible (Fig. 3). We also repeated these analyses separately for di*ff*erent brain networks. We found that networks traditionally associated with higher-level cognitive functions tended to provide more informative brain patterns than networks traditionally associated with lower level cognitive functions (Fig. 4). Finally, we examined the most dominant components of the brain activity patterns from each experimental condition. We used a reverse inference approach (Rubin et al., 2017) to identify the terms in the neuroimaging literature most commonly associated with the corresponding maps. As summarized in Figure 6E, we found that the intact and paragraph conditions tended to weight higher-level cognitive processes more than lower-level cognitive processes, whereas the word condition weighted lower-level processes more than higher-level processes and the rest condition showed no reliable di*ff*erence in high-level versus low-level weightings. Taken together, our findings indicate that the informativeness and compressibility of our brain activity patterns are task-dependent, and these properties change systematically with factors like cognitive richness and depth of processing.

Our explorations of informativeness and compressibility are related to a much broader literature on the correlational and causal structure of brain activity patterns and networks (Adachi et al., 2012; Bassett & Sporns, 2017; Brovelli et al., 2004; Bullmore & Sporns, 2009; Dhamala et al., 2008; Korzeniewska et al., 2008; Lynn & Bassett, 2021; Owen et al., 2021; Preti et al., 2017; Rogers et al., 2007; Rubinov & Sporns, 2010; Sizemore et al., 2018; Smith, Beckmann, et al., 2013; Smith, Vidaurre, et al., 2013; Sporns & Betzel, 2016; Sporns & Honey, 2006; Sporns & Zwi, 2004; Srinivasan et al., 2007; Tomasi & Volkow, 2011; Yeo et al., 2011). Correlations or causal associations between di*ff*erent brain regions simultaneously imply that full-brain activity patterns will be compressible and also that those activity patterns will contain redundancies. For example, the extent to which activity patterns at one brain area can be inferred or predicted from activity patterns at other areas (e.g., Owen et al., 2020; Scangos et al., 2021), reflects overlap in the information available in or represented by those brain areas. If brain patterns in one area are recoverable using brain patterns in another area, then a “signal” used to convey the activity patterns could be compressed by removing the recoverable activity. Predictable (and therefore redundant) brain activity patterns are also more robust to signal corruption. For example, even if the activity patterns at one region are unreadable or unreliable at a given moment, that unreliability could be compensated for by other regions’ activity patterns that were predictive of the unreliable region. Whereas compressible brain patterns are robust to spatial signal corruption, high versus low informativeness reflects a similar (though dissociable; e.g., Fig. 1) tradeo*ff* between expressiveness and robustness of *temporal* patterns. Highly informative brain patterns (by our measure; i.e., patterns that yield greater temporal decoding accuracy) are expressive about ongoing experiences or cognitive states, since each moment’s pattern is reliably distinguishable from other moments’ patterns. However, when each moment’s pattern is unique, brain activity becomes less robust to temporal signal corruption. Our finding that brain activity patterns becomes more informative (i.e., less robust to temporal signal corruption) and compressible (i.e., more robust to spatial signal corruption) when cognitive engagement is higher suggests that our brain may optimize its activity patterns to prioritize either temporal or spatial robustness, according to task demands.

Our findings that informativeness and compressibility change with task demands may follow from task-dependent changes in full-brain correlation patterns. A number of prior studies have found that so-called “functional connectivity” (i.e., correlational) patterns vary reliably across tasks, events, and situations (Cole et al., 2014; Owen et al., 2021; Simony et al., 2016; Smith et al., 2009). By examining how these task-dependent changes in correlations a*ff*ect informativeness and compressibility, our work suggests a potential reason why the statistical structure of brain activity patterns might vary with cognitive task or with cognitive demands. For lower-level tasks, or for tasks that require relatively little “deep” cognitive processing, our brains may optimize activity patterns for robustness and redundancy over expressiveness, for example to maximize reliability. For higher-level tasks, or for tasks that require deeper cognitive processing, our brains may sacrifice some redundancy in favor of greater expressiveness.

One potential limitation of our work concerns how our measure of informativeness might generalize across di*ff*erent tasks, cognitive representations, and processes. Our use of across-participant temporal decoding accuracy as a proxy for informativeness is motivated in part by prior work that introduced across-participant similarity (in time-varying response to a stimulus) as a means of identifying stimulus-driven brain activity patterns (Simony et al., 2016). Intuitively, only activity patterns that are driven by the stimulus would be expected to synchronize (i.e., be time-locked to the stimulus) across participants. This approach implicitly removes idiosyncratic responses (e.g., neural patterns that are *not* similar across people). However, there are also some published examples, including in our own prior work, that indicate that some types of stimulus-evoked activity will be missed by across-participant comparisons. For example we have reported how brain regions like the ventromedial prefrontal cortex (vmPFC) show stimulus-driven responses that are, for the most part, *not* similar across people. In that paper (and drawing on other work), we suggest that the vmPFC seems to represent or support highly idiosyncratic internal states, like a*ff*ective responses. Although we would consider the vmPFC to be a “high-level” region (e.g., we consider a*ff*ect to be a relatively high-level aspect of cognition), the measure of informativeness that we used in our current study would identify regions like the vmPFC as having *low* informativeness. This is because across-participant decoding accuracy (our proxy measure for informativeness) will only be high for representations or responses that are common across people.

Relatedly, even in the experimental conditions we describe as “less cognitively engaging,” we think it likely that high-level thought or cognitive processing is still present. Rather, we suggest that these high-level representations will tend to be more idiosyncratic when the stimulus is less engaging, and therefore less constraining on people’s thoughts. Nonetheless, even during highly engaging tasks, people may engage in idiosyncratic stimulus-driven processes. For example, people might retrieve personal information as they listened to the story. Those retrievals could happen at di*ff*erent times for di*ff*erent people according to each individual’s prior experiences. Even when those sorts of retrievals happen to be temporally synchronized across people, the specific memories or information being retrieved might still be idiosyncratic. Our measure of informativeness is insensitive to these processes. Further, even in response to an identical stimulus, task instructions or participants’ internal goals could change the relationship between compressibility and informativeness. Some work has shown that the “dimensionality” of neural representations can change systematically with task complexity, even in response to an identical stimulus (Mack et al., 2020). Taken together, we expect that the way we have defined informativeness in this paper, and the specific dataset we examined, are likely to have influenced our findings. While we see our approach as a reasonable first step, we also suggest that future work should explore alternative measures of informativeness and compressibility, and should examine how these measures vary across di*ff*erent tasks and datasets.

In the information theory sense (Shannon, 1948), when a signal is transmitted using a fixed alphabet of “symbols,” the information rate decreases as the signal is compressed (e.g., fewer symbols transmitted per unit time, using an alphabet with fewer symbols, etc.). Our finding that each individual brain component (symbol) becomes more informative as cognitive richness increases suggests that the “alphabet” of brain activity patterns is also task-dependent. Other work suggests that the representations that are *reflected* by brain activity patterns may also change with task demands. For example, our brains may represent the same perceptual stimulus di*ff*erently depending on which aspects of the stimulus or which combinations of features are task-relevant (Mack et al., 2020).

We found that di*ff*erent brain networks varied in how informative and compressible their activity patterns were across experimental conditions (e.g., Fig. 4). This might follow from evolutionary optimizations that reflect the relevant constraints or demands placed on those networks. One possibility is that cortex is organized in a hierarchy of networks “concerned with” or selective to di*ff*erent levels of processing or function. To the extent that di*ff*erent levels of processing (e.g., low-level sensory processing versus “deeper” higher-level processing) reflect di*ff*erent stimulus timescales (e.g., Manning, 2023), the network di*ff*erences we observed might also relate to the timescales at which each network is maximally sensitive (Baldassano et al., 2017; Hasson et al., 2008; Lerner et al., 2011; Regev et al., 2018).

Our reverse inference analyses (Figs. 5, 6) also provide some insights into how neural activity patterns change with cognitive engagement or task demands. Prior work has shown that the components and network “parcels” identified through covarying activity patterns can be highly similar even across di*ff*erent tasks (including “rest,” e.g., Laird et al., 2011; Smith et al., 2009). We replicated this basic finding in that the first principal components from all four experimental conditions were strikingly similar (e.g., see the leftmost columns of Figs. 5A and 6C, D). We also found some small, though statistically reliable, systematic changes in the weights associated with di*ff*erent cognitive functions across conditions (Fig. 6E). This result provided an additional way of characterizing network-level di*ff*erences across conditions (Fig. 4E). Taken together, these findings suggest that although similar networks may be involved in di*ff*erent tasks, the ways in which those networks are engaged may vary systematically with task demands.

### Concluding remarks

Cognitive neuroscientists are still grappling with basic questions about the fundamental “rules” describing how our brains respond, and about how brain activity patterns and the associated underlying cognitive representations and computations are linked. We identified two aspects of brain activity patterns, informativeness and compressibility, that appear to change systematically with task demands and across brain networks. We speculate that these changes may reflect ongoing tradeo*ff*s between how robust to signal corruption versus how expressive about ongoing cognitive states our brains’ activity patterns are. Our work also provides a new framework for evaluating these tradeo*ff*s in other datasets, or in future studies.

## Methods

We measured properties of recorded neuroimaging data under di*ff*erent task conditions that varied systematically in cognitive engagement and depth of processing. We were especially interested in how *informative* and *compressible* the activity patterns were under these di*ff*erent conditions (Fig. 1). Unless otherwise noted, all statistical tests are two-sided, all error bars and error ribbons denote bootstrap-estimated 95% confidence intervals across participants, and all reported *p*-values are corrected for multiple comparisons using the Benjamini-Hochberg procedure (Benjamini & Hochberg, 1995).

### Functional neuroimaging data collected during story listening

We examined an fMRI dataset collected by Simony et al. (2016) that the authors have made publicly available at arks.princeton.edu*/*ark:*/*88435*/*dsp015d86p269k. The dataset comprises neuroimaging data collected as participants listened to an audio recording of a story (intact condition; 36 participants), listened to temporally scrambled recordings of the same story (17 participants in the paragraph-scrambled condition listened to the paragraphs in a randomized order and 36 in the word-scrambled condition listened to the words in a randomized order), or lay resting with their eyes open in the scanner (rest condition; 36 participants). Full neuroimaging details may be found in the original paper for which the data were collected (Simony et al., 2016). Procedures were approved by the Princeton University Committee on Activities Involving Human Subjects, and by the Western Institutional Review Board (Puyallup, WA). All subjects were native English speakers with normal hearing and provided written informed consent. We have excerpted the relevant portions of the dataset documentation here to provide information about the scanning parameters and preprocessing steps used to generate the data we analyzed (the original descriptions may be found at the above link):

> Subjects were scanned in a 3T full-body MRI scanner (Skyra; Siemens) with a sixteen-channel head coil. For functional scans, images were acquired using a T2* weighted echo planer imaging (EPI) pulse sequence [repetition time (TR), 1500 ms; echo time (TE), 28 ms; flip angle, 64^◦^], each volume comprising 27 slices of 4 mm thickness with 0 mm gap; slice acquisition order was interleaved. In-plane resolution was 3 × 3 mm^2^ [field of view (FOV), 192 × 192 mm^2^]. Anatomical images were acquired using a T1-weighted magnetization-prepared rapid-acquisition gradient echo (MPRAGE) pulse sequence (TR, 2300 ms; TE, 3.08 ms; flip angle 9^◦^; 0.89 mm^3^ resolution; FOV, 256 mm^2^). To minimize head movement, subjects’ heads were stabilized with foam padding. Stimuli were presented using the Psychophysics toolbox (Brainard, 1997; Pelli, 1997). Subjects were provided with an MRI compatible in-ear mono earbuds (Sensimetrics Model S14), which provided the same audio input to each ear. MRI-safe passive noise-canceling headphones were placed over the earbuds, for noise removal and safety.
>
> Functional data were preprocessed and analyzed using FSL (www.fmrib.ox.ac.uk/fsl), including correction for head motion and slice-acquisition time, spatial smoothing (6 mm FWHM Gaussian kernel), and high-pass temporal filtering (140 s period). Preprocessed data were aligned to coplanar and high-resolution anatomicals and the standard MNI152 brain, and interpolated to 3-mm isotropic voxels.

The intact and word conditions each comprised 300 TRs (7.5 minutes) per participant. The paragraph condition comprised 272 TRs (6.8 minutes) per participant. The rest condition comprised 400 TRs (10 minutes) per participant.

### Hierarchical topographic factor analysis (HTFA)

Following our prior related work, we used HTFA (Manning et al., 2018) to derive a compact representation of the neuroimaging data. In brief, this approach approximates the timeseries of voxel activations (44,415 voxels) using a much smaller number of radial basis function (RBF) nodes (in this case, 700 nodes, as determined by an optimization procedure; Manning et al., 2018). This provides a convenient representation for examining full-brain activity patterns and network dynamics. All of the analyses we carried out on the neuroimaging dataset were performed in this lower-dimensional space. In other words, each participant’s data matrix was a number-of-timepoints (*T*) by 700 matrix of HTFA-derived factor weights (where the row and column labels were matched across participants). Code for carrying out HTFA on fMRI data may be found as part of the BrainIAK toolbox (Capota et al., 2017; Kumar et al., 2021), which may be downloaded at brainiak.org.

We also considered alternative approaches to obtaining compact representations of the neuroimaging data, including network parcellations (e.g., Gordon et al., 2016; Schaefer et al., 2018). Whereas network parcellations are typically derived from large resting state datasets, HTFA may be applied to much smaller datasets. In our prior work, we showed that HTFA applied to the same dataset used here can explain full-brain activity to within a maximum of 0.25 standard deviations of each voxel’s observed activity in the original dataset, taken across all voxels, images, and participants, using the 700-node representation we also employed here (Manning et al., 2018). Some of the explanatory power of HTFA comes from the fact that each node’s influence falls o*ff* smoothly with distance to its center. Intuitively, the result is a representation that looks like a lightly spatially smoothed version of the original data, but where the degree of smoothing varies across the brain according to how spatially autocorrelated the local activity patterns are.

### Network permutation tests

In our analyses of how informativeness varied across brain networks (Fig. 4), we considered the possibility that the correlations with network order might be influenced by the numbers of nodes in each network. We designed a permutation-based procedure to address this possibility, whereby we repeated the above analyses using shu*ffl*ed network labels. Specifically, for each of *n*_1_ *=* 10 iterations, we randomly shu*ffl*ed (without replacement) the network labels of the HTFA nodes, and then we re-ran our entire decoding analysis pipeline, including applying PCA with 3…*m* features for each condition (where *m* is the number of nodes in the given network), and then running 100 cross-validation runs of the decoding procedure for each condition and number of components. This resulted in 10 sets of shu*ffl*ed data, where each network had the same numbers of nodes, but where the decoding results no longer maintained the fidelity of each individual network.

We sampled the original and shu*ffl*ed datasets (with replacement) to create *n*_2_ *=* 1000 bootstrap samples. For each bootstrap sample, we computed the correlations between the decoding accuracies and network order for each condition and number of components. This yielded a distribution of *n*_2_ correlation values for each condition, for both the original and shu*ffl*ed datasets. We then compared the distributions of Spearman’s ρ values for the original and shu*ffl*ed datasets using two-sided independent samples Welch’s *t*-tests.

### Principal components analysis (PCA)

We applied group PCA (Smith et al., 2014) separately to the HTFA-derived representations of the data (i.e., factor loadings) from each experimental condition. Specifically, for each condition, we considered the set of all participants’ *T* by 700 factor weight matrices. We used group PCA to project these 700-dimensional matrices into a series of shared *k*-dimensional spaces, for *k* ∈ {3, 4, 5, …, 700}. This yielded a set of number-of-participants matrices, each with *T* rows and *k* columns.

### Temporal decoding

We sought to identify neural patterns that reflected participants’ ongoing cognitive processing of incoming stimulus information. As reviewed by Simony et al. (2016), one way of homing in on these stimulus-driven neural patterns is to compare activity patterns across individuals. In particular, neural patterns will be similar across individuals to the extent that the neural patterns under consideration are stimulus-driven, and to the extent that the corresponding cognitive representations are reflected in similar spatial patterns across people (Simony & Chang, 2020). Following this logic, we used an across-participant temporal decoding test developed by Manning et al. (2018) to assess the degree to which di*ff*erent neural patterns reflected ongoing stimulus-driven cognitive processing across people. The approach entails using a subset of the data to train a classifier to decode stimulus timepoints (i.e., moments in the story participants listened to) from neural patterns. We use decoding (forward inference) accuracy on held-out data, from held-out participants, as a proxy for the extent to which the inputted neural patterns reflected stimulus-driven cognitive processing in a similar way across individuals.

#### Forward inference and decoding accuracy

We used an across-participant correlation-based classifier to decode which stimulus timepoint matched each timepoint’s neural pattern. For a given value of *k* (i.e., number of principal components), we first used group PCA to project the data from each condition into a shared *k*-dimensional space. Next, we divided the participants into two groups: a template group, G_template_ (i.e., training data), and a to-be-decoded group, G_decode_ (i.e., test data). We averaged the projected data within each group to obtain a single *T* by *k* matrix for each group. Next, we correlated the rows of the two averaged matrices to form a *T* by *T* decoding matrix, Λ. In this way, the rows of Λ reflected timepoints from the template group, while the columns reflected timepoints from the to-be-decoded group. We used Λ to assign temporal labels to each timepoint (row) from the test group’s matrix, using the row of the training group’s matrix with which it was most highly correlated. We repeated this decoding procedure, but using G_decode_ as the template group and G_template_ as the to-be-decoded group. Given the true timepoint labels (for each group), we defined the decoding accuracy as the average proportion of correctly decoded timepoints, across both groups (where chance perfomance is 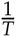). In Figures 2 and 3 we report the decoding accuracy for each condition and value of *k*, averaged across *n =* 100 cross validation folds.

### Reverse inference

To help interpret the brain activity patterns we found within the contexts of other studies, we created summary maps of each principal component, for each experimental condition. Each principal component comprises 700 “weights” on each of the HTFA-derived RBF nodes (see *Hierarchical Topographic Factor Analysis*). For each node, we evaluated its RBF at the locations of every voxel in the standard 2 mm MNI152 template brain and multiplied the RBF by the node’s weight. The sum of these weighted RBF activation maps provides a full-brain image, in MNI152 space, of the given principal component (Fig. S3).

Next, we considered 80 topics estimated using Latent Dirichlet Allocation (Blei et al., 2003) applied to 9,204 functional neuroimaging articles in the Neurosynth database (Rubin et al., 2017). The topics, as well as associated brain maps identified using Neurosynth, were identified and reported in several prior studies (Chen et al., 2020; Fox et al., 2014; Sul et al., 2017). The topic labels for each topic were generated automatically with the following ChatGPT (OpenAI, 2023) prompt: “Please help me come up with intuitive labels for topics I found by fitting a topic model to thousands of neuroscience and psychology articles. I’ll paste in the top 10 highest-weighted words for each topic, and I’d like you to respond with a suggested label. For each topic, please respond with just the topic label and no other formatting or text. Here are the next topic’s top words:” followed by a comma-separated list of the given topic’s top-weighted words reflected in the Table S1. For some topics, ChatGPT responded with a longer-form response rather than a concise topic label. In these instances, on a case-by-case basis, we used a second follow-up prompt to achieve the given topic’s label: “Could you please come up with a more concise label for that topic?”. We then manually identified a set of 11 cognitive labels that were intended to encapsulate a representative range of widely studied low-level and high-level cognitive functions. In choosing the set of cognitive labels, we jointly considered each topic’s ChatGPT-derived topic label, along with the top-weighted words for the topic. We attempted to generate a concise set of labels that still spanned the full set of cognitive functions reflected across the 80 topics. Topics that appeared unrelated to specific cognitive functions (e.g., topics related to specific methods or clinical themes) are designated with dashes in Table S1.

Finally, following an approach used in several prior studies (Chen et al., 2020; Fox et al., 2014; Sul et al., 2017) we treated the correlation between a given component’s brain map and each topic’s brain map as an approximate measure of how much the component was reflective of the given topic. This resulted in a set of 80 “weights” (correlation coe*ffi*cients) for each component’s brain map, with one weight per Neurosynth-derived topic.

### Ranking cognitive processes

We manually identified 11 cognitive labels spanning the set of 80 Neurosynth-derived topics: cognitive control, language processing, memory, emotion, social cognition, spatial cognition, attention, reward, sensory perception, motor control, and resting state. We then used ChatGPT to automatically “rank” the processes from high-level to low-level using the following prompt: “Please rank these cognitive processes from highest-level to lowest-level, where higher values indicate higher-order or higher-level processes. Return the result as a csv file with a header row and two columns: ‘Cognitive label’ and ‘Rank’. Here are the processes: cognitive control, language processing, memory, emotion, social cognition, spatial cognition, attention, reward, sensory perception, motor control, resting state”. Table S2 displays the output.

We recognize that ChatGPT is not omniscient, nor should it be treated as an expert cognitive neuroscientist. We therefore reviewed ChatGPT’s responses carefully by hand to verify that they seemed reasonable to us. Whereas prior work has often constructed such rankings by hand, we see our use of ChatGPT in this case as a small additional “sanity check” on our rankings that helped us to be slightly more objective than if we had simply created the rankings ourselves manually.

In the analysis presented in Figure 6E, we summarize di*ff*erence in topic weightings across experimental conditions. In particular, we sought to characterize how the dominant neural patterns evoked by each experimental condition weighted on di*ff*erent cognitive functions. For each of the top five principal components from each experimental condition (Fig. 5), we computed the average weights for each of the 11 manually identified (and ChatGPT-ranked) cognitive labels described above (Tab. S2). We then fit a line separately for each experiment condition (*x*-values: cognitive rank; *y*-values: weights). In carrying out this analysis, we used a bootstrap procedure to estimate the variability in the slopes of the regression lines, whereby we repeated this process for each of *n =* 100 iterations, each time resampling (with replacement) the set of observed ranks and weights. This procedure yielded distributions of 100 estimated slopes for each experimental condition. We used these distributions to compare the slopes across experimental conditions and to estimate 95% confidence intervals.

### Synthetic data

To help illustrate the relationship between informativeness and compressibility (Fig. 1), we generated four synthetic datasets, varying in informativeness and compressibility. Each dataset comprised simulated observations of *K =* 25 features across *N =* 100 timepoints, from each of *S =* 10 participants. To create each dataset, we first constructed a “template” matrix of *N* timepoints by *K* features. We then generated participant-specific data by adding independent noise to each entry in template matrix, drawn from the unit normal distribution (i.e., with a mean of 0 and a variance of 1). We repeated this process for each participant, yielding *S* participant-specific matrices for each dataset.

Since we estimate informativeness using the temporal decoding accuracy across participants, highly informative data will tend to have observations that are highly timepoint specific. Relatively uninformative data, in contrast, will tend to have more similar observations across timepoints. To generate data with “high informativeness,” we constructed template matrices whose rows (observations) were drawn independently from zero-mean multivariate normal distributions. The covariances of these distributions were determined according to the desired compressibility of the data, as described below. We used a multi-step process to generate data with “low informativeness.” First we generated new template matrices using the same procedure as for the “high informativeness” datasets. We then multiplied each matrix by a constant (ρ *=* 0.1) and computed the cumulative sum of each matrix’s rows. This yielded matrices whose rows were highly similar across observations.

Compressibility reflects the extent to which decoding accuracy is a*ff*ected by reducing the number of components used to represent the data. Highly compressible data will tend to exhibit more similarities across features, whereas less compressible data will tend to show greater independence across features. To generate data with “high compressibility,” we set the covariance matrix of the multivariate normal distribution to a toeplitz matrix whose first row was given by [*K*, *K* − 1, …, 1]. To generate data with “low compressibility,” we set the covariance matrix to the identity matrix.

Template matrices for datasets with high informativeness and high compressibility, high informativeness and low compressibility, low informativeness and high compressibility, and low informativeness and low compressibility are displayed in Figure 1C. The corresponding decoding curves are displayed in Figure 1D.

## Data and code availability

All of the code used to produce the figures and results in this manuscript, along with links to the corresponding data, may be found at github.com*/*ContextLab*/*pca paper.

## Acknowledgements

We acknowledge discussions with Rick Betzel, Luke Chang, Emily Finn, and Jim Haxby. Our work was supported in part by NSF CAREER Award Number 2145172 to J.R.M. The content is solely the responsibility of the authors and does not necessarily represent the o*ffi*cial views of our supporting organizations. The funders had no role in study design, data collection and analysis, decision to publish, or preparation of the manuscript.

## Author contributions

Conceptualization: J.R.M. and L.L.W.O. Methodology: J.R.M. and L.L.W.O. Implementation: J.R.M. and L.L.W.O. Analysis: J.R.M. and L.L.W.O. Writing, Reviewing, and Editing: J.R.M. and L.L.W.O. Funding acquisition: J.R.M. Supervision: J.R.M.

