## Supplemental Materials for "High-level cognition is supported by information-rich but compressible brain activity patterns"

Lucy L. W. Owen[1, 2] and Jeremy R. Manning[1, *]
[1]Department of Psychological and Brain Sciences,
Dartmouth College, Hanover, NH
[2]Carney Institute for Brain Sciences,
Brown University, Providence, RI

December 26, 2023

| Topic label | Cognitive label | Term 1 | Term 2 | Term 3 | Term 4 | Term 5 | Term 6 | Term 7 | Term 8 | Term 9 | Term 10 |
|---|---|---|---|---|---|---|---|---|---|---|---|
| Cognitive control and task performance | Cognitive control | tasks | control | network | conditions | comparison | performed | common | correlates | experiment | pre |
| Developmental aging and maturation | - | age | adults | children | development | adolescents | aging | developmental | adolescence | childhood | adult |
| Eye movements and visual attention | Attention | eye | gaze | eyes | visual | saccades | movements | saccade | direction | gait | target |
| Facial and voice recognition | Sensory perception | recognition | familiar | identity | unfamiliar | voice | familiarity | route | voices | fg | facial |
| Social interaction and contextual behavior | Social cognition | context | game | human | interaction | ppi | contextual | contexts | agency | interactions | partner |
| Language processing and semantic knowledge | Language processing | semantic | words | word | lexical | verbal | language | tasks | naming | fluency | phonological |
| Experimental design and behavioral performance | - | trials | stimulus | responses | trial | reaction | time | event | target | events | times |
| Genetic polymorphisms and risk factors | - | carriers | allele | gene | genotype | met | genetic | polymorphism | val | comt | rs |
| Sensorimotor integration and movement control | Motor control | motor | movement | movements | sensorimotor | primary | finger | control | imagery | sensory | execution |
| Drug addiction and substance abuse | - | cocaine | users | drug | bpd | controls | cannabis | addiction | craving | dependent | heroin |
| Music perception and auditory processing | Sensory perception | music | musical | pitch | auditory | musicians | sequences | rhythm | listening | beat | singing |
| Menstrual cycle and hormonal regulation | - | phase | women | cycle | phases | menstrual | hf | expression | sex | luteal | follicular |
| Cognitive functions and role playing | Cognitive control | role | play | human | plays | cognitive | evidence | critical | distinct | key | subregions |
| Inhibition and gender differences | - | inhibition | women | inhibitory | sex | females | gender | males | stop | male | female |
| Somatosensory stimulation and motor control | Motor control | stimulation | somatosensory | tms | tactile | primary | touch | motor | rtms | transcranial | sensory |
| Multisensory integration and perception | Sensory perception | auditory | visual | sensory | modality | sounds | integration | sound | stimulus | primary | modalities |
| Social perception and empathy | Social cognition | social | empathy | experience | people | person | responses | perspective | individuals | attachment | empathic |
| Gesture recognition and visual attention | Attention | target | gestures | targets | orientation | visual | distractors | gesture | grasp | distractor | location |
| Experimental design | - | design | block | blocks | event | mixed | condition | ca | writing | blocked | runs |
| Alcohol cue reactivity | Reward | cues | cue | alcohol | anticipation | cued | preparation | anticipatory | exposure | expectancy | preparatory |
| Neuroimaging and metabolism | - | pet | tomography | emission | positron | flow | glucose | binding | metabolism | metabolic | receptor |
| Abnormalities in schizophrenia | - | schizophrenia | controls | reduced | abnormalities | symptoms | deficits | matched | control | abnormal | schizophrenic |
| Eating and body weight | - | food | taste | body | weight | eating | women | obese | reward | foods | caloric |
| Sleep and olfactory processing | Sensory perception | sleep | olfactory | odor | sd | deprivation | odors | rem | wakefulness | night | wake |
| Alzheimer's disease and mild cognitive impairment | - | ad | disease | mci | alzheimer | cognitive | impairment | mild | amci | controls | atrophy |
| Working memory and executive function | Memory | memory | load | tasks | verbal | maintenance | performance | term | difficulty | dual | cognitive |
| Moral decision making and phobias | - | moral | phobia | phobic | ec | guilt | spider | decision | art | phobics | intentional |
| Language laterality | Language processing | language | asymmetry | organization | human | dominance | asymmetries | lateralized | handed | located | representation |
| Attention | Attention | attention | attentional | visual | spatial | top | selective | stimulus | control | orienting | target |
| Resting-state brain activity in smokers | Resting state | reho | smokers | resting | smoking | nicotine | homogeneity | sci | controls | rs | spontaneous |
| Social cognition/judgment | Social cognition | social | judgments | mind | mental | theory | mentalizing | cognition | judgment | person | people |
| Reward and decision making | Reward | reward | decision | choice | outcomes | rewards | monetary | decisions | anticipation | responses | outcome |
| ADHD and attention deficits | Attention | adhd | disorder | attention | deficit | children | hyperactivity | deficits | control | controls | reduced |
| Neurobiological variability and individual diff... | - | individual | relationship | local | dependent | change | global | responses | neuronal | magnitude | lower |
| Spatial cognition | Spatial cognition | spatial | space | location | locations | navigation | virtual | visual | visuospatial | visuo | position |
| Therapeutic interventions and training | - | training | acupuncture | therapy | cr | control | trained | improvement | mindfulness | stimulation | induced |
| Color perception and deception | Sensory perception | color | search | feature | deception | features | responses | colour | dimension | lying | conjunction |
| Neurodegenerative diseases and disorders | - | disease | pd | controls | atrophy | clinical | motor | dementia | sd | multiple | sclerosis |
| Cognitive control and interference | Cognitive control | conflict | control | interference | stroop | incongruent | selection | congruent | trials | cognitive | monitoring |
| Structural MRI and brain volume analysis | - | volume | gray | voxel | gm | morphometry | grey | volumes | vbm | density | age |
| Fear conditioning and extinction | Emotion | fear | switching | conditioning | responses | stimulus | extinction | cs | switch | threat | conditioned |
| Skill learning and expertise | Memory | learning | practice | learned | sequence | performance | training | sequences | skill | implicit | motor |
| PTSD and trauma | Emotion | ptsd | trauma | stress | disorder | traumatic | posttraumatic | childhood | survivors | exposure | controls |
| Neural oscillations and electrophysiology | - | frequency | source | alpha | amplitude | beta | gamma | recorded | frequencies | potential | simultaneous |
| Temporal dynamics of stimulus processing | Sensory perception | time | sustained | duration | onset | period | stage | timing | delay | transient | event |
| Tinnitus and hearing loss | Sensory perception | tinnitus | loss | hearing | status | driving | subjective | objective | unfair | offers | rejection |
| Abstract categories and representations | Language processing | category | adaptation | representations | categorization | categories | abstract | stimulus | representation | features | knowledge |
| Pain perception and sensory stimulation | Sensory perception | pain | painful | stimulation | somatosensory | intensity | noxious | heat | chronic | sensory | nociceptive |
| Body and primates | - | body | human | humans | monkeys | itch | primates | species | monkey | bodies | macaque |
| Phonological processing in reading | Language processing | reading | chinese | phonological | visual | language | readers | dyslexia | characters | children | word |
| Rule-based performance and complexity | Cognitive control | rule | complexity | size | force | rules | effect | grip | amount | distance | artificial |
| Autism Spectrum Disorder (ASD) and social impai... | Social cognition | asd | autism | social | spectrum | individuals | controls | disorders | children | td | reduced |
| Major depression disorder and emotions | Emotion | depression | mdd | depressed | depressive | major | mood | disorder | sad | controls | control |
| Blindness and vision | Sensory perception | blind | visual | sighted | individuals | humor | laughter | congenitally | blindness | plasticity | braille |
| Deafness and sign language | Language processing | condition | conditions | deaf | hearing | sign | signs | referential | signers | ci | nss |
| Genetic risk and familial factors in psychosis | - | risk | genetic | sz | siblings | relatives | individuals | unaffected | factors | family | psychosis |
| Action observation and imitation | Motor control | action | actions | observation | motor | mirror | goal | imitation | execution | directed | movements |
| Cognitive performance and control | Cognitive control | performance | cognitive | control | memory | executive | test | tasks | individuals | behavioral | deficits |
| Mental disorders and controls | - | disorder | ocd | bipolar | hc | bd | controls | ts | control | abnormalities | compulsive |
| Pharmacological effects of placebo and drug adm... | - | placebo | dopamine | effect | drug | ht | administration | blind | dopaminergic | double | mg |
| Personality and anxiety | Social cognition | anxiety | personality | trait | individuals | scores | traits | threat | disorders | social | avoidance |
| Mental imagery and math abilities | Cognitive control | mental | imagery | numerical | arithmetic | rotation | calculation | magnitude | digit | tasks | mathematical |
| Priming and repetition effect | Memory | priming | repetition | suppression | repeated | effect | implicit | literal | target | prime | metaphors |
| Working memory and error monitoring | Memory | wm | error | errors | prediction | performance | correct | monitoring | ltm | feedback | predicted |
| Sentence comprehension and syntax | Language processing | sentences | language | sentence | comprehension | syntactic | semantic | verbs | verb | word | meaning |
| Resting state networks | Resting state | network | resting | default | mode | rest | intrinsic | cognitive | correlations | seed | spontaneous |
| Episodic memory encoding and retrieval | Memory | memory | encoding | retrieval | recognition | episodic | items | successful | subsequent | recollection | recall |
| Visual object recognition | Sensory perception | object | visual | objects | shape | images | scenes | scene | selective | recognition | stream |
| Effective causal modeling of neural networks | - | effective | causal | modeling | dynamic | network | top | influence | modulation | interactions | causality |
| Relational reasoning and fluid intelligence | - | reasoning | FALSE | TRUE | intelligence | relational | belief | relations | nonverbal | fluid | analogical |
| Lesion and stroke rehabilitation | - | lesions | lesion | stroke | damage | injury | controls | recovery | patient | tracts | integrity |
| Affective valence and feedback processing | - | negative | positive | feedback | valence | arousal | bias | positively | negatively | evaluation | swallowing |
| Autobiographical memory in epilepsy | Memory | autobiographical | events | future | epilepsy | episodic | memory | memories | past | mtle | personal |
| Evidence and effect in behavioral studies | - | evidence | provide | effect | behavioral | underlying | demonstrated | understanding | potential | responses | support |

| Topic label | Cognitive label | Term 1 | Term 2 | Term 3 | Term 4 | Term 5 | Term 6 | Term 7 | Term 8 | Term 9 | Term 10 |
|---|---|---|---|---|---|---|---|---|---|---|---|
| Stress and physiological responses | - | stress | tdcs | cortisol | autonomic | heart | responses | rate | regulation | physiological | induced |
| Speech and language processing | Language processing | speech | language | auditory | production | perception | comprehension | listening | acoustic | linguistic | prosody |
| Network interactions and evidence in human systems | - | network | evidence | human | systems | support | process | distinct | integration | provide | engaged |
| Neuroimaging techniques | - | standard | images | individual | time | image | voxel | spatial | test | clinical | mapping |
| Visual perception of motion and form | Sensory perception | motion | visual | perception | perceptual | biological | dynamic | moving | human | static | illusion |
| Emotional processing and regulation | Emotion | emotional | emotion | neutral | facial | expressions | affective | responses | negative | regulation | emotions |

Table S1: **Neurosynth-derived topics.** We report the top-weighted terms for each of 80 topics identified using Latent Dirichlet Allocation (Blei et al., 2003) applied to 9,204 functional neuroimaging articles in the Neurosynth database (Rubin et al., 2017). See *Reverse inference* for additional information.

| Cognitive label | Rank |
|---|---|
| Cognitive control | 10 |
| Language processing | 9 |
| Memory | 8 |
| Emotion | 7 |
| Social cognition | 6 |
| Spatial cognition | 5 |
| Attention | 4 |
| Reward | 3 |
| Sensory perception | 2 |
| Motor control | 1 |
| Resting state | 0 |

Table S2: **Ranking cognitive processes.** The table displays the output of a ChatGPT (OpenAI, 2023) prompt asking for a ranking of the cognitive processes reflected in the labels from Table S1. See *Ranking cognitive processes* for additional detail.

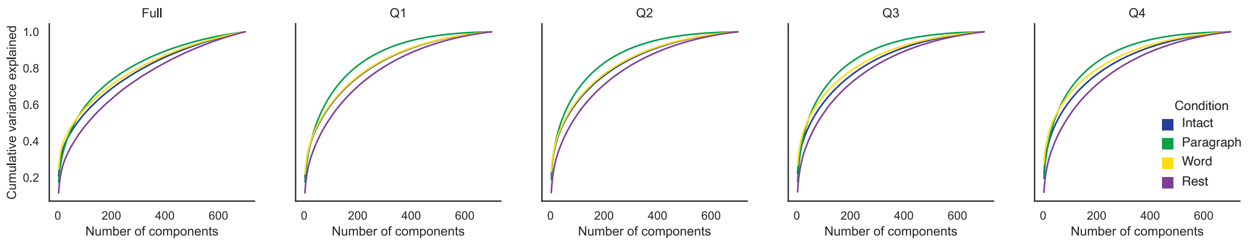


Figure S1: **Cumulative variance explained by component, condition, and part.** Each panel displays the cumulative variance explained in the neuroimaging data as a function of the number of principal components. Colors denote experimental conditions. The left panel displays results for all data, and the right panels display results separated by story segment (Q1: first quarter; Q2: second quarter; Q3: third quarter; Q4: fourth quarter).

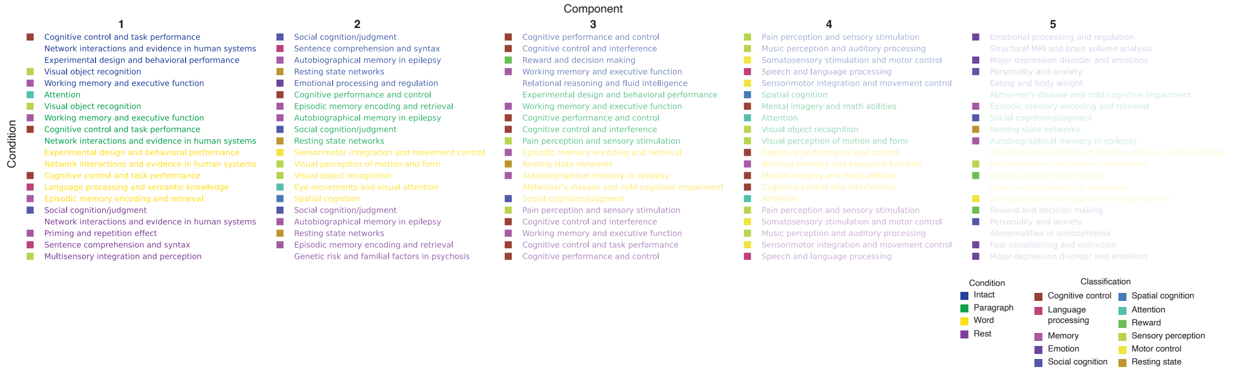


Figure S2: **Highest-weighted topics associated with the highest-weighted components by condition, broken down by story segment.** Each group of five rows corresponds to an experimental condition (denoted by color, as indicated in the legend in the lower right), and the columns and shading correspond to the component number (ranked by proportion of variance explained). The colored squares in front of many of the topics denote manually identified cognitive labels (Tab. S1).

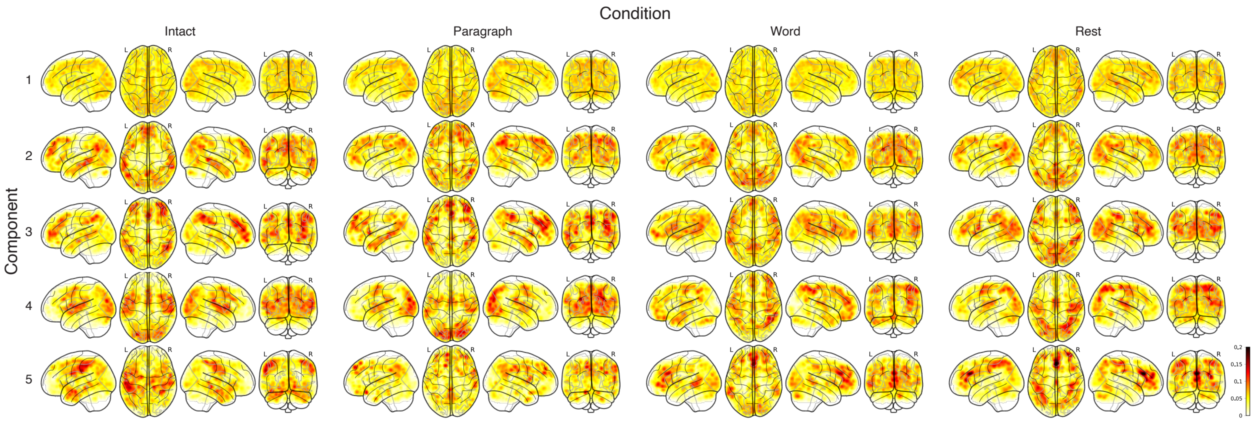


Figure S3: **Brain maps by component and condition.** For the top five highest-weighted principal components (rows), from each experimental condition (columns), the components' brain maps are projected onto four views: left sagittal, axial, right sagittal, and coronal. The color scale is the same for all panels and matches the coloring in Figure 5C.

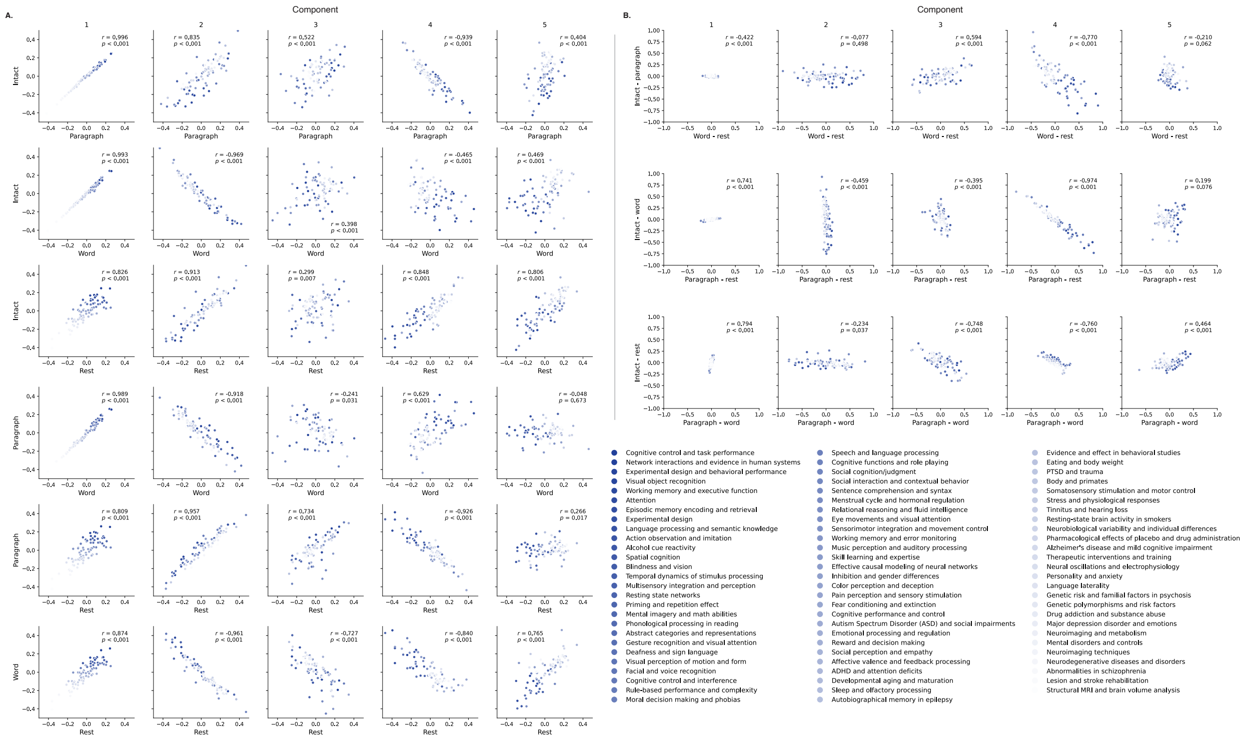


Figure S4: **Comparisons between per-component topic correlations across conditions.** Each sub-panel displays a scatterplot comparing the per-topic correlations for two or more experimental conditions. Each dot denotes the correlations for a single topic (indicated by the legend on the right). The topics are colored according to the ranked order of the correlations between the topic's brain maps and the brain map for the first principal component in the intact condition. **A. Comparisons between correlations for each pair of experimental conditions.** The conditions being compared are marked on the $x$ and $y$ axes. Each sub-panel (column) reflects the correlations for one principal component. **B. Comparisons between *differences* in correlations for pairs of experimental conditions.** In these sub-panels, the $x$ and $y$ coordinates reflect differences in correlations for the indicated experimental conditions, for the given component (column). All panels: the across-topic correlations reported in each panel are between each topic's $x$ and $y$ coordinates.

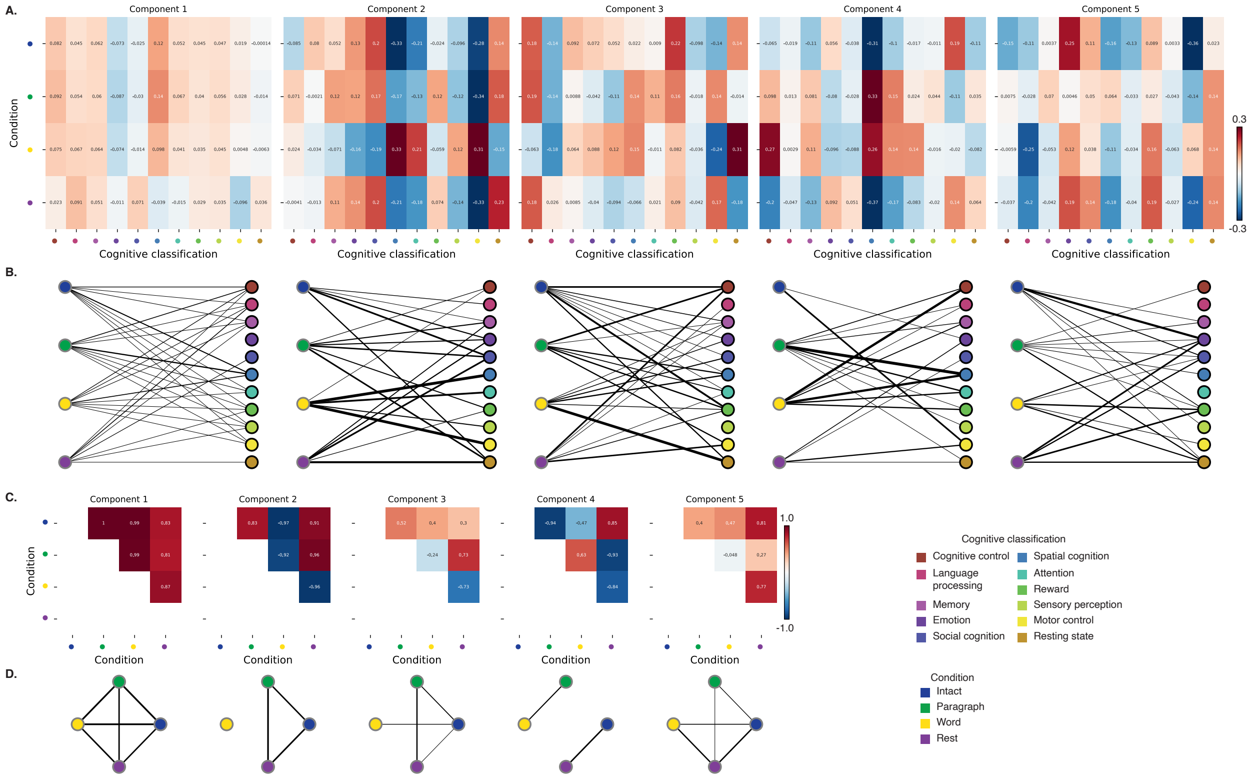


Figure S5: **Functions associated with top-weighted components by condition. A. Top-weighted topics by condition.** Here we display per-condition (rows, indicated by colored dots) topic correlations, averaged across topics that pertain to each of several broad cognitive functions (columns within each sub-panel, indicated by colored dots). Each sub-panel reflects correlations for the components indicated in the panel titles. A legend for the condition and cognitive function classifications is displayed in the lower right of the figure. Table S1 provides a list of each topic's top-weighted terms, along with each topic's manually labeled cognitive classification. A full list of the topics most highly associated with each component may be found in Figure S2. **B. Associations between per-condition components and cognitive functions.** The network plots denote positive average correlations between the component images for each condition (gray-outlined dots on the left sides of each network; colors denote conditions) and topic-specific brain maps associated with each indicated cognitive function (black-outlined dots on the right sides of each network; colors denote cognitive functions). The line thicknesses are proportional to the correlation values (correlation coefficients are noted in the heat maps in Panel A). **C. Correlations between each principal component, by condition.** The heat maps display the correlations between the brain maps (Fig. S3) for each principal component (sub-panel), across each pair of conditions (rows and columns of each sub-panel's matrix, indicated by colored dots). **D. Associations between per-condition topic weights, by component.** Each sub-panel's network plot summarizes the pattern of correlations between the topic correlations from each of the $n^{\text{th}}$ top-weighted principal components (sub-panel), for each experimental condition (gray-outlined dots). The line thicknesses are proportional to the correlation values (correlation coefficients are noted in the heat maps in Panel C).
